# Microbial communities thriving in agave fermentations are locally influenced across diverse biogeographic regions

**DOI:** 10.1101/2024.03.22.586289

**Authors:** Angélica Jara-Servin, Luis D. Alcaraz, Sabino I. Juarez-Serrano, Aarón Espinosa-Jaime, Ivan Barajas, Lucia Morales, Alexander DeLuna, Antonio Hernández-López, Eugenio Mancera

## Abstract

The production of traditional agave spirits in Mexico is a deeply rooted traditional process that relies on environmental microorganisms to ferment the cooked must from agave plants. Analysis of these microorganisms provides the opportunity to understand the dynamics of the microbial communities in the interface of natural and human-associated environments in a biologically and culturally rich region of the world. Here, we performed 16S and ITS amplicon sequencing of close to 100 fermentation tanks from 42 distilleries throughout Mexico. The *Agave* species used, production practices, climatic conditions, and biogeographic characteristics varied considerably among sites. Yet, we did find taxa present in most fermentations suggesting that there is a core of microorganisms that are hallmarks of these communities. These core taxa are represented by hundreds of OTUs showing large intra-specific variation. The only variable that was consistently associated with the composition of both bacterial and fungal communities was the distillery, suggesting that microbial composition is determined by the local production practices and unique attributes of each site. Fermentation stage, climate and producing region were also associated with the community composition, but only for prokaryotes. Analysis of microbial composition in several tanks within three distilleries also revealed taxa that were enriched in specific fermentation stages or agave species. Our work provides a comprehensive analysis of the microbiome of agave fermentations, contributing key knowledge for its management and conservation.

## 1 INTRODUCTION

Mexico stands out for its biological and cultural diversity, a result of its intricate geological and social history. For example, it has one of the world’s highest numbers of reptile, oak, and pine species (Farjon 1996, Nixon 2006, Suazo-Ortuño et al. 2023). Additionally, it is one of the most multilingual regions, and its population is predominantly admixed (Eberhard et al. 2020, Sohail et al. 2023). However, we know relatively less about its microbial diversity in both natural and human-associated habitats. The production of spirits from agave plants is a highly ingrained activity in Mexico. It is yet unclear whether this practice dates to pre-Columbian times, but currently it is performed from the southernmost provinces of Mexico to the northern states that border with the USA (Serra Puche and Lazcano Arce 2016, Cabrera-Toledo et al. 2022). Its widespread distribution is probably related to the abundance of agave plants throughout the Mexican territory as this region is the center of diversity of the genus (Trejo et al. 2018).

*Tequila* is arguably the best known and most commercial of the agave spirits, nowadays produced in highly industrial settings. However, most other agave distillates, such as *mezcal*, *raicilla* or *bacanora*, are produced in a more artisanal way that is characterized by open, “spontaneous” fermentations in which the producers do not rely on an established inoculum of microorganisms (Arellano-Plaza et al. 2022). Instead, the cooked and macerated agave hearts are fermented by microorganisms coming from the surrounding environment. Therefore, the characterization of the microbiome involved in agave fermentations is not only essential to better understand the production of these beverages of traditional and commercial importance, but it also provides insight into the natural communities of microorganisms in Mexico.

The vast area where agave spirits are produced — a region that is larger in size than Western Europe — shows a wide range of environmental conditions and production practices (Arellano-Plaza et al. 2022). For instance, there are distilleries located almost at sea level, while others stand at 2,000 meters above it, encompassing arid, semi-arid and subhumid climates in subtropical and tropical areas. This results in a wide range of temperatures and annual rainfalls determining, among other things, the agave species that can grow in each region. Over 50 different species of agave have been reported to be used for the production of agave spirits and their sugar, nitrogen and other metabolite content could vary among them (Colunga-GarciaMarin et al. 2017). The cooking process hydrolyses fructans, the most abundant carbohydrates in agave plants, releasing free sugars such as fructose, for the fermentation step. However, cooking also produces compounds that can inhibit microorganism growth such as 5-hydroxymethylfurfural and furfural (Mancilla-Margalli and López 2002). Together with metabolites from the agave plants such as saponins, these compounds can make agave fermentations a challenging environment for microorganisms. For an in-depth review of agave fermentations as an ecological environment for microorganisms see Colón-González M., *et al*. (2024).

The most common microorganisms identified in agave fermentations using culture-based microbiological methods, with their intrinsic biases, are ascomycetous yeasts and lactic acid bacteria (Colón-González M., *et al*. 2024, Escalante-Minakata et al. 2008, Kirchmayr et al. 2017, Gallegos-Casillas et al. 2023). Previous studies have mainly focused on yeasts since they are thought to be the main contributors to ethanol production. The most extensive characterization to date of the fungal communities of traditional agave fermentations recently revealed a core of six ascomycetous yeasts that were commonly isolated from these fermentation (Gallegos-Casillas et al. 2023). These species had also been isolated in previous studies, even when they were only carried out in a handful of factories mostly located in the state of Oaxaca, the main mezcal producing state (Kirchmayr et al. 2017, Nolasco-Cancino et al. 2018). The different species of yeast involved in these fermentations are not only thought to contribute to ethanol production, but also to the synthesis of other compounds with organoleptic properties.

Much less is known about the prokaryotic microbiome of agave fermentations and bacteria in other anthropogenic fermentations are often considered contaminants that can lead to spoilage. The three previous studies that did analyze prokaryotes in traditional fermentations for the production of agave spirits, isolated mainly species of lactic acid bacteria, but also acetic acid bacteria and spore-forming bacteria (Escalante-Minakata et al. 2008, Narvaez-Zapata et al. 2010, Kirchmayr et al. 2017). The ethanol producing capabilities of some of these species suggested that bacteria could actually have a more important role than previously thought in the production of traditional agave spirits (Kirchmayr et al. 2017). Despite the previous efforts using culture-based microbiological methods, there are still many basic open questions regarding the microbiome of traditional agave fermentations. In fact, it is unclear whether the isolated yeast species are indeed the predominant fungi in these fermentations. It is possible that other fungi are also abundant but are not commonly isolated with classical microbiological methods. The types of bacteria that play a role in the different regions where agave spirits are produced are also unknown.

To get deeper insights into both the bacterial and fungal microbiome of agave fermentations we performed 16S rRNA gene (16S) and ribosomal internal transcribed spacer (ITS) amplicon sequencing from must samples collected from 99 fermentation tanks from 42 different distilleries throughout the regions where agave spirits are produced in Mexico. Our results showed that despite the diversity in production practices and biogeographical parameters there is a core of species that defines these microbial communities throughout the country. Many of these taxons coincide with species previously isolated from these fermentations, but we also identified prevalent species that had never been associated with agave fermentations before. Although Ascomycota were by far the most common fungi, we also identified fungi from five other phyla, showing overall considerable fungal diversity. The bacterial microbiome is even more diverse than the fungal component, and it is dominated by species of lactic acid bacteria. The only association of the composition of the microbiome that we observed in both bacteria and fungi was with the distillery to which the samples belong. To our knowledge, our work represents the first countrywide analysis employing metagenomic approaches of the microbiome of the fermentations used for traditional agave spirit production, considerably expanding our understanding of these communities.

## 2 MATERIALS AND METHODS

### 2.1 Collection of agave fermentation samples, DNA extraction and sequencing

Agave fermentation samples were taken as part of the field work performed by the YeastGenomesMx consortium throughout the seven regions where agave spirits are produced in Mexico. As described before (Gallegos-Casillas et al. 2023), for each sample, 4 ml of agave ferment were taken with a sterile serological pipet at arm’s reach from each of the 99 sampled tanks and were stored in cryogenic vials to be immediately frozen in liquid nitrogen. Associated metadata (Supplementary Table 1) was collected in the field for each sample as previously described (Gallegos-Casillas et al. 2023). At the laboratory, samples were stored at -80 °C until processing. Total DNA was extracted using the ZymoBIOMICS DNA Miniprep Kit (D4300) from Zymo Research. Extractions were performed from 500 μl of sample following the manufacturer protocol but adding a 15 minute incubation at 70 °C after lysis and before centrifugation and filtration. DNA quantity, purity and integrity were monitored with a Nanodrop, fluorometry in a Qubit and by performing agarose gel electrophoresis. 16S rRNA gene and ITS library preparation and sequencing were performed by Zymo Research in an Illumina MiSeq platform and generating a minimum of 25,000 PE 300bp reads per amplicon type for each sample. The 16s V3 and V4 regions were sequenced for samples from distillery D1 used to assess technical and biological replicability, while only the V4 region was sequenced for the rest of the samples. pPNA and mPNA blockers were used to avoid plant DNA amplification and sequencing.

### 2.2 16S rRNA gene and ITS region amplicons sequence processing

Access to the detailed protocol and bioinformatic methods used to process and analyze 16S rRNA genes and ITS amplicon sequences can be found on GitHub (/github.com/ajaraservin/mezcal). Amplicon data from the pulque microbiome were obtained from publicly available raw reads (Rocha-Arriaga et al. 2020). Quality assessment was performed on all 16S rRNA gene amplicon libraries using FastQC (https://www.bioinformat ics.babraham.ac.uk/projects/fastqc/). The CASPER v0.8.2 assembler was used to merge the pair-end reads (Kwon et al. 2014). To process all samples in the same manner, only the V4 region was employed, removing the V3 sequence from the D1 distillery and from the pulque data (Rocha-Arriaga et al. 2020). QIIME’s identify_chimeric_seqs.py using ChimeraSlayer (Caporaso et al. 2010) script with BLAST fragments was used to identify chimeric sequences. All samples were concatenated and clustered into Operational Taxonomic Units (OTUs) using a 97% identity threshold with cd-hit-est (Li and Godzik 2006). Taxonomy assignment was performed using QIIME’s scripts (Caporaso et al. 2010) against the Silva v 138 database (Yilmaz et al., 2014). After singleton, chimera and contamination screening, the remaining sequences were aligned, and a phylogenetic tree was constructed using FastTreeMP v2.1.11 (Price et al. 2009).

ITS pair-end reads were merged using the CASPER assembler and then subjected to a quality check using fastq_quality_filter (Q<20) from the FASTX-Toolkit (http://hannonlab.cshl.edu/fastx_toolkit/). After assembly with PANDASEQ v2.11 and clustering, the all-eukaryotes UNITE v8.3 database (Nilsson et al. 2019) was used for taxonomic assignment up to the species level. All chimeras and non-fungi sequences were discarded using parallel_identify_chimeric_seqs.py and filter_otus_from_otu_table.py scripts (Caporaso et al. 2010).

### 2.3 Diversity and statistical analysis

OTUs were used to analyze both the α and β-diversity of our agave spirit and pulque samples. Phyloseq (McMurdie and Holmes 2013), ggplot2 (Wickham 2016), vegan (Oksanen et al. 2022), and R default packages (Team 2021) were used for the analysis. Observed, Chao1, Shannon, and Simpson diversity indices were calculated to assess α-diversity, while taxonomic abundance profiles were obtained through clustering at different taxonomic levels. Dendrograms were constructed using the hclust method. Constrained analysis of principal coordinates (CAP) on an unweighted UniFrac matrix (Lozupone and Knight 2005) of 16S ribosomal RNA (16S rRNA) gene sequences was used to evaluate ß-diversity according to the different variables. Similarly, a Jaccard similarity matrix (Knight et al. 2018) of ITS sequences was used for the PcoA ordination method. Both matrices were evaluated using the ANOSIM statistical function (Varsos et al. 2016). Differential OTUs abundance was estimated using R’s package DESeq2 (Love et al. 2014). Finally, Geographic Distance Matrix Generator v.1.2.3 (Ersts) was used to generate a geographic distance matrix for all agave spirit samples. This geographic distance matrix was used to perform a Mantel test (Xia and Sun 2017), using an unweighted UniFrac distance matrix for bacteria and a Jaccard similarity matrix for fungi. Detailed bioinformatic and statistical methods are available at GitHub (/github.com/ajaraservin/mezcal).

## 3 RESULTS

### 3.1 A countrywide collection of agave musts from diverse biogeographical regions

At the time of writing, the YeastGenomesMx consortium had sampled over 90 distilleries in all the regions where agave spirits are produced in Mexico. For the characterization of the microbiome of agave fermentations described here we focused on 99 fermentation tanks from 42 distilleries, 67 of which used *A. angustifolia*, the most used species for traditional agave spirits. *A. tequilana*, the agave species used to produce tequila, is by far the most cultivated agave species, however, tequila production is mostly industrial, involving inoculation of the fermentations with commercial yeast strains and was therefore not considered in this work. Focusing the analysis on distilleries that use *A. angustifolia* allowed us to test the impact of other variables besides agave species on the microbiome of the fermentations, mainly whether there is an association between the location of the distillery and the microbiome. Given that there are regions where *A. angustifolia* is not employed, we also included fermentations of other eight agave species as well as mixtures of two or three species for the production of blended spirits. The species used in these other distilleries are *A. americana, A. durangensis, A. inaequidens, A. karawinskii, A. mapisaga, A. potatorum, A. rhodacantha,* and *A. salmiana.* Using this subset that includes fermentations of a variety of agaves we were also able to test the possible effect of the plant substrate on microbial diversity.

The 42 distilleries are located in 31 different municipalities of the states of Durango, Guanajuato, Michoacan, Oaxaca, Puebla, Tamaulipas, Jalisco and Sonora. In the six first states, the spirit produced is known as *mezcal* while in Jalisco is known as *raicilla* and in Sonora as *bacanora*. In addition, several producers prefer to use the general term “*destilado de agave*” (Spanish for agave spirit) to name their spirits. All three first spirits have their own designation of origin, although the overall production process is similar. The two more distant distilleries that were sampled are over 2,000 km apart in a straight line, from 29°52’ North and 109°33’ West to 16° North and 96°31’ West. The altitude of the distilleries also varied considerably, from below 500 meters above sea level to over 2,500 meters, and covered climates from tropical to semi-desertic. Figure 1 shows the geographic location of the distilleries analyzed in this study and some of their geographical characteristics.

**Figure 1.**
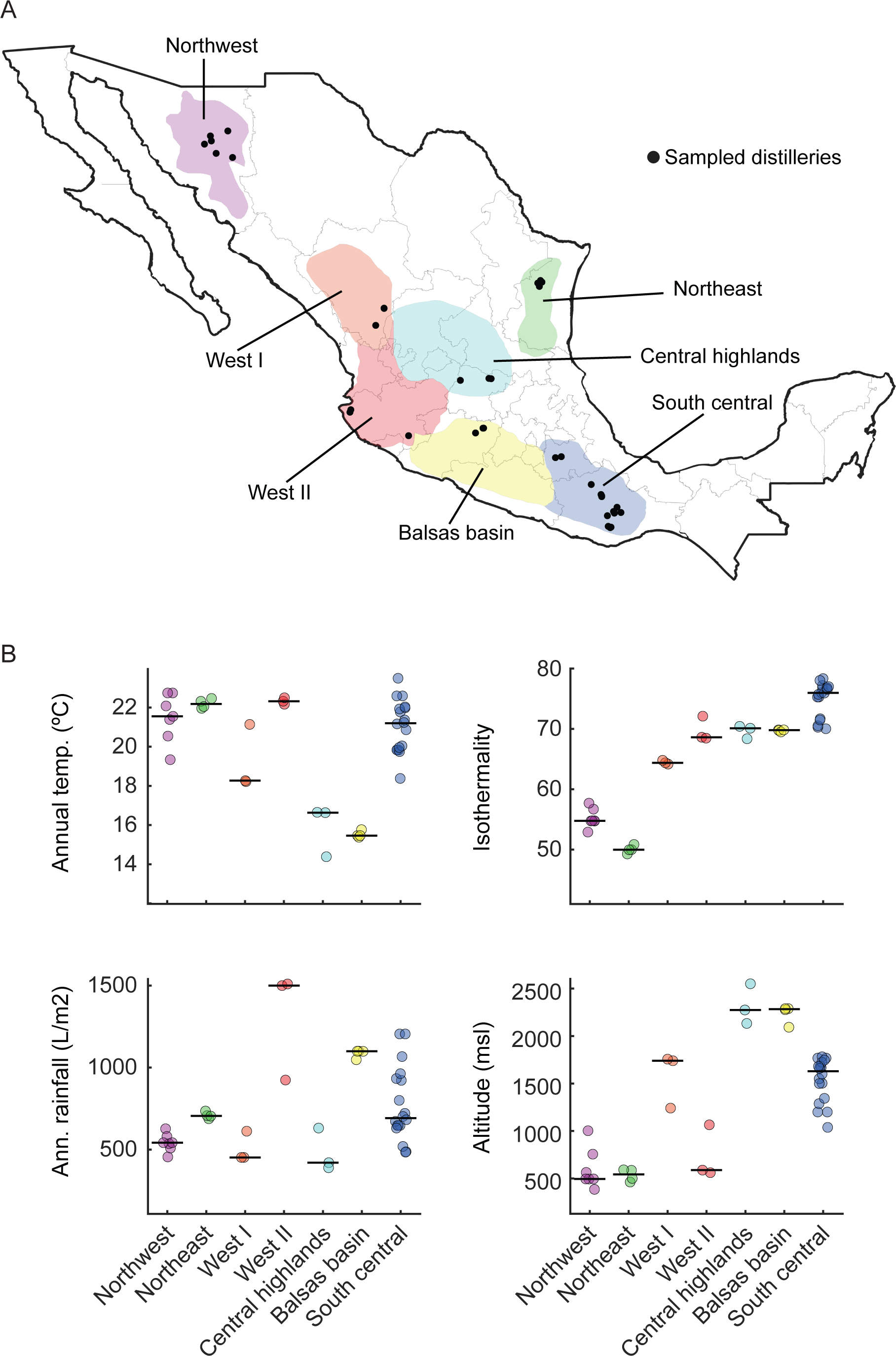
Distribution and characteristics of the agave spirit distilleries. A) Map of Mexico showing the geographical location of the 42 distilleries (black dots) from which the 99 fermentation tanks were sampled in the corresponding producing region which are indicated as polygons in different background colors. Producing regions were defined according to (Aguirre 2006) mostly based on the agave species and production practices employed. The West I and West II regions were considered a single region (West) in the original publication, but after consulting with the authors, the region was divided to take into account the differences that we observed in the production practices of the distilleries of those areas (Gallegos-Casillas et al. 2023). B) Climatic characteristics of the locations of the agave fermentation tanks. Top left) average annual temperature, top right) isothermality, bottom right) average annual rainfall, and bottom left) height above mean sea level.

All the distilleries that were included in this study but three, used buried stone or brick ovens and the cooked agave hearts were grinded either manually, using mills powered by animals or employing motorized mills. The tanks sampled spanned fermentation times from the day the fermentation was formulated to full maturity when the must was being transferred to the alembic for distillation. In terms of fermentation days, we sampled tanks that had less than a day to those that had been fermenting for close to two months. The material of the tanks was diverse including wood, plastic, clay, steel, cement, and even cattle hide, but wood was by far the most common followed by plastic among the distilleries sampled. The specific characteristics of the distilleries and fermentation tanks of the sequenced samples are detailed in Supplementary Table 1.

### 3.2 Bacterial diversity in agave fermentations across Mexico

In this study, we assessed and compared the bacterial community of agave fermentations throughout Mexico by 16S metagenomic sequencing. Of the 99 sequenced samples, only 91 were sequenced successfully, from which we generated a total of 9,622,960 paired-end 16S reads. From these, 5,191,846 assembled sequences passed the quality control parameters with a mean of 53,524.18 ± 21,879.91 sequences per sample. Using a 97% identity cut-off we obtained 10,094 OTUs, clustered into 888 genera that matched a total of 29 distinct prokaryotic phyla. On average, each sample contained a richness of 1,002.08 ± 316.47 OTUs. The average expected Chao1 richness was 1,690.17 ± 500.67 OTUs which showed that our effort covered a major part of the expected bacterial OTUs. The average Shannon-Weiner (H’) and Simpson’s (D) diversity indices across samples were 3.26 ± 0.66 and 0.86 ± 0.12, respectively.

Throughout the samples, the composition of the prokaryotic communities at the phylum level was similar, and was predominantly composed by Firmicutes (71.82%), Proteobacteria (14%), Actinobacteriota (4.82%), Bacteroidota (4.02%), Verrucomicrobiota (0.76%), and Patescibacteria (0.65%). From the 71 identified bacterial classes, Bacilli dominated (6,802 OTUs; 68.33%) followed far away by Alphaproteobacteria (971 OTUs; 9.75%), Gammaproteobacteria (971 OTUs; 9.75%), Actinobacteria (412 OTUs; 4.13%), Bacteroidia (399 OTUs; 4.00%), Clostridia (373 OTUs; 3.74%), and Parcubacteria (46 OTUs; 0.46%). The ten most abundant genera were *Weissella* (948 OTUs; 739,215 reads), *Paucilactobacillus* (258 OTUs; 637,238 reads), *Lentilactobacillus* (749 OTUs; 600,744 reads), *Leuconostoc* (678 OTUs; 464,976 reads), *Oenococcus* (266 OTUs; 335,694 reads), *Lactiplantibacillus* (509 OTUs; 270,402 reads), *Liquorilactobacillus* (373 OTUs; 270,842 reads), *Lacticaseibacillus* (587 OTUs; 233,680 reads), *Acetobacter* (236 OTUs; 224,959 reads) and *Secundilactobacillu*s (572 OTUs; 255,747 reads). All of them, but *Acetobacter,* are lactic acid bacteria. We also observed considerable diversity within the identified genera. For example, more than 60% of all genera were represented by more than one OTU and over 10% by ten or more (Supplementary Table 2).

The most abundant genera described above, except *Acetobacter,* were present in all the distilleries that we sampled. Overall, there were 15 genera present throughout the sites analyzed, all lactic acid bacteria, that could be considered the core bacterial microbiome of agave fermentations (Table 1). If a lenient definition of the core is used, in addition to these 15, there were 11 other genera that were present in 80% or more of the distilleries and in all the seven agave-spirit producing regions (Table 1). By far, the majority (73%) of these 26 genera were lactic acid bacteria. The taxonomic abundance profile of prokaryotes is shown in Figure 2 and the complete list of genera is provided in Supplementary Table 2.

**Table 1.**
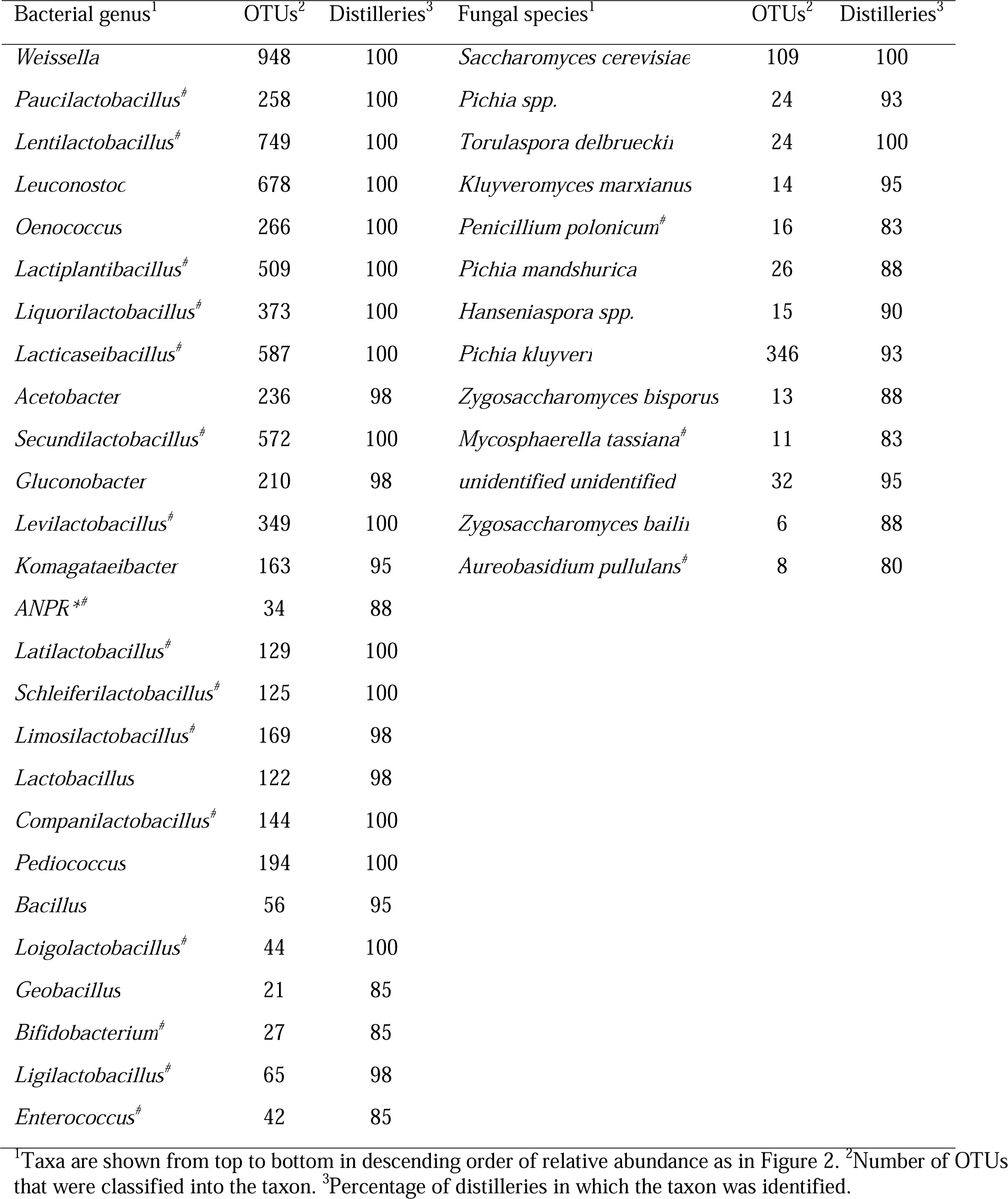
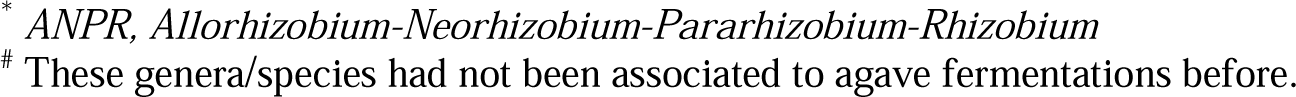
Bacterial and fungal core microbiome of traditional agave fermentations.

| Bacterial genus<sup>1</sup> | OTUs<sup>2</sup> | Distilleries<sup>3</sup> | Fungal species<sup>1</sup> | OTUs<sup>2</sup> | Distilleries<sup>3</sup> |
| --- | --- | --- | --- | --- | --- |
| *Weissella* | 948 | 100 | *Saccharomyces cerevisiae* | 109 | 100 |
| *Paucilactobacillus*<sup>#</sup> | 258 | 100 | *Pichia spp.* | 24 | 93 |
| *Lentilactobacillus*<sup>#</sup> | 749 | 100 | *Torulaspora delbrueckii* | 24 | 100 |
| *Leuconostoc* | 678 | 100 | *Kluyveromyces marxianus* | 14 | 95 |
| *Oenococcus* | 266 | 100 | *Penicillium polonicum*<sup>#</sup> | 16 | 83 |
| *Lactiplantibacillus*<sup>#</sup> | 509 | 100 | *Pichia mandshurica* | 26 | 88 |
| *Liquorilactobacillus*<sup>#</sup> | 373 | 100 | *Hanseniaspora spp.* | 15 | 90 |
| *Lacticaseibacillus*<sup>#</sup> | 587 | 100 | *Pichia kluyveri* | 346 | 93 |
| *Acetobacter* | 236 | 98 | *Zygosaccharomyces bisporus* | 13 | 88 |
| *Secundilactobacillus*<sup>#</sup> | 572 | 100 | *Mycosphaerella tassiana*<sup>#</sup> | 11 | 83 |
| *Gluconobacter* | 210 | 98 | *unidentified unidentified* | 32 | 95 |
| *Levilactobacillus*<sup>#</sup> | 349 | 100 | *Zygosaccharomyces bailii* | 6 | 88 |
| *Komagataeibacter* | 163 | 95 | *Aureobasidium pullulans*<sup>#</sup> | 8 | 80 |
| *ANPR*<sup>*#</sup> | 34 | 88 |  |  |  |
| *Latilactobacillus*<sup>#</sup> | 129 | 100 |  |  |  |
| *Schleiferilactobacillus*<sup>#</sup> | 125 | 100 |  |  |  |
| *Limosilactobacillus*<sup>#</sup> | 169 | 98 |  |  |  |
| *Lactobacillus* | 122 | 98 |  |  |  |
| *Companilactobacillus*<sup>#</sup> | 144 | 100 |  |  |  |
| *Pediococcus* | 194 | 100 |  |  |  |
| *Bacillus* | 56 | 95 |  |  |  |
| *Loigolactobacillus*<sup>#</sup> | 44 | 100 |  |  |  |
| *Geobacillus* | 21 | 85 |  |  |  |
| *Bifidobacterium*<sup>#</sup> | 27 | 85 |  |  |  |
| *Ligilactobacillus*<sup>#</sup> | 65 | 98 |  |  |  |
| *Enterococcus*<sup>#</sup> | 42 | 85 |  |  |  |

<sup>1</sup>Taxa are shown from top to bottom in descending order of relative abundance as in Figure 2. <sup>2</sup>Number of OTUs that were classified into the taxon. <sup>3</sup>Percentage of distilleries in which the taxon was identified.
\* ANPR, *Allorhizobium-Neorhizobium-Pararhizobium-Rhizobium*

**Figure 2.**
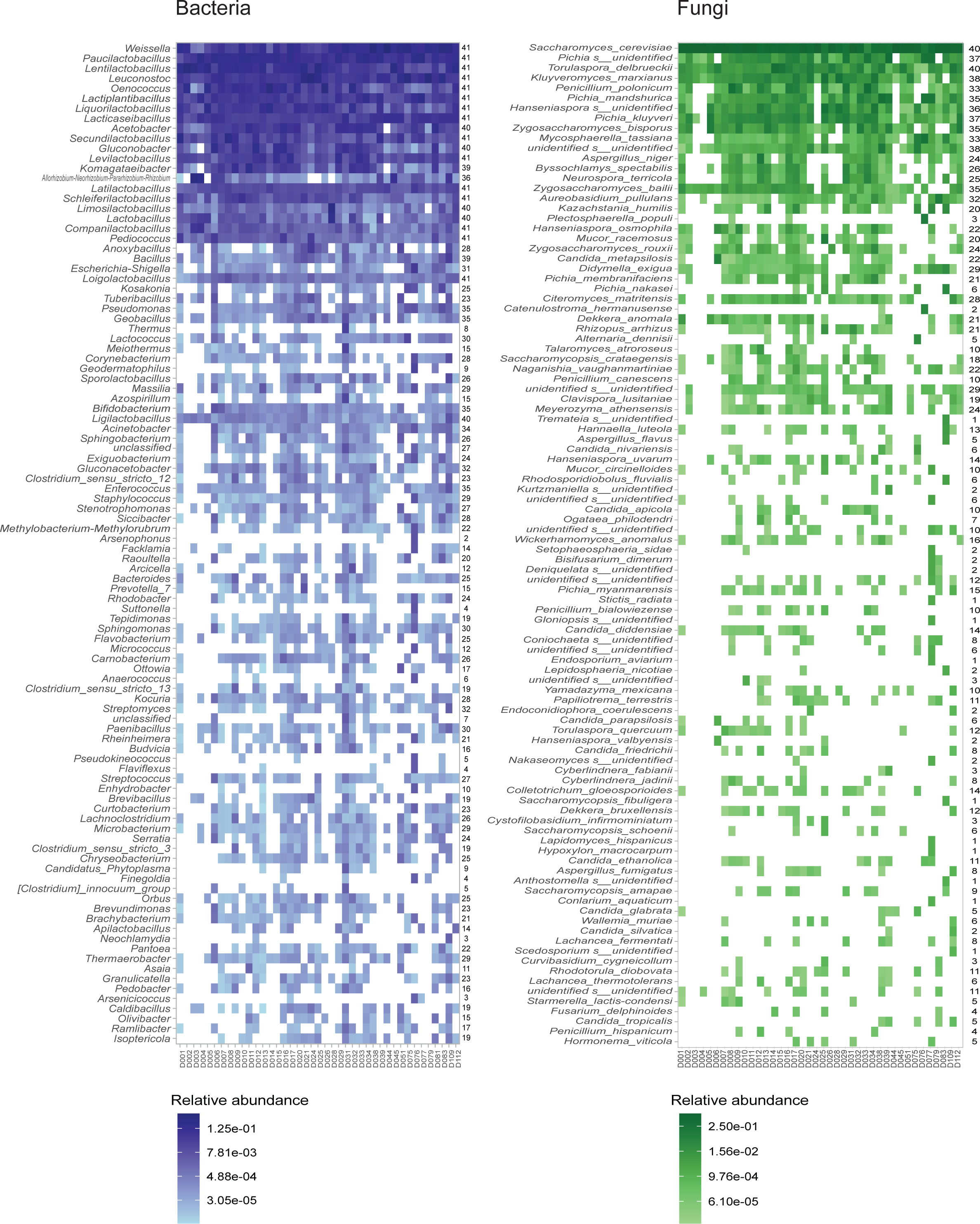
Bacterial and fungal composition of the microbial communities of agave fermentations. Heatmaps showing the relative abundance of the bacterial genera (left panel) and fungal species (right panel) identified in each of the distilleries. Only the 100 most abundant taxa are shown ordered from top to bottom according to their overall relative abundance. Numbers at the right of each heatmap show the total number of distilleries in which the taxon was identified. All the bacterial and fungal taxa identified are reported in Supplementary Table 2.

From the 784 bacterial genera that could be classified, close to 98% had not been previously associated with traditional agave fermentations using culture-based microbiological methods (Colón-González M., *et al*., 2024). From the core bacterial microbiome, 16 (61.5%) genera had not been reported in previous literature (Table 1). It is important to note that some of these genera were recently defined which could explain why they had not been previously reported in these fermentations. Furthermore, from the 20 genera that had been isolated from these fermentations in the past, our survey only missed *Zymomonas*. However, *Zymomonas* is not present as a genus in the most current 16S reference database in which the sequences that belonged to this genus have been reassigned to five genera of the family Sphingomonadaceae, four of which we did identify in agave fermentations. Overall, these results showed that our sampling effort contributed considerably to the understanding of the prokaryotic composition of traditional agave fermentations, a part of this microbiome that had been mostly neglected.

### 3.3 Fungal diversity in agave fermentations across Mexico

To characterize the fungal microbiome of agave fermentations we performed amplicon sequencing of the ITS region from the must samples. A total of 87 samples were successfully sequenced yielding a total of 5,244,553 paired-end ITS reads. From these, 2,212,845 assembled sequences passed the quality control parameters with an average of 47,656.9 ± 1,025.3 reads per sample. By clustering the sequences at a phylogenetic distance of 3%, we identified a total of 1,118 ITS OTUs. An average of 92.85 ± 39.72 OTUs were observed in the 87 samples, while the average expected richness (Chao1) was 153.92 ± 70.44. Therefore, as with prokaryotes, our effort covered most of the expected diversity per sample. The diversity in each sample was considerably lower than for bacteria with an average Shannon-Weiner diversity index (H’) of 1.20 ± 0.74 and an average Simpson diversity index (D) of 0.44 ± 0.27. By far, most of the fungal OTUs belonged to the phylum Ascomycota (1,035 OTUs; 92.57%), but we also identified Basidiomycota (40 OTUs; 3.57%), Mucoromycota (7 OTUs 0.62%), Chytridiomycota (2 OTUs; 0.17%), Mortierellomycota (1 OTU; 0.08%) and Rozellomycota (1 OTU; 0.08%). 11 fungal OTUs (0.98%) remained unidentified even at the level of phyla.

The 1,118 identified OTUs were distributed in 325 fungal species. The ten most abundant fungal species were *Saccharomyces cerevisiae* (109 OTUs; 1,461,038 reads), *Pichia mandshurica* (26 OTUs; 1,000,851 reads), an unidentified species of *Pichia* (24 OTUs; 100,495 reads), *Torulaspora delbrueckii* (24 OTUs; 78,018 reads), *Penicillium polonicum* (16 OTUs; 59,716 reads), *Kluyveromyces marxianus* (14 OTUs; 54,326 reads), *Zygosaccharomyces bisporus* (13 OTUs; 46,565 reads), an unidentified species of *Hanseniaspora* (15 OTUs; 44,753 reads), *Pichia kluyveri* (346 OTUs; 43,204 reads), and another unidentified species (32 OTUs; 25,695 reads). These species accounted for 94.26% of the total number of reads. Surprisingly, *P. polonicum* and *Mycosphaerella tassiana* (11 OTUs; 24,623 reads) had, to our knowledge, never been associated with agave fermentations before. As for many prokaryote genera, large intra-specific diversity was observed; 81 (23.5%) species had more than two OTUs and 19 (5.5%) more than ten (Supplementary Table 2).

Only *S. cerevisiae* and *T. delbrueckii* were present in all the distilleries, but there were other eleven fungal species identified in 80% or more of the sites and in all the seven producing regions. These 13 species could be considered the core fungal microbiome of agave fermentations (Table 1). One of these core species could not be identified even at the level of phylum, but the rest are all ascomycetes. Numerous rare species were also detected, with 125 found in only one fermentation tank, and we also identified 35 and 34 fungal OTUs that could not be assigned to any species or genus, respectively. The taxonomic abundance profile of fungi is shown in Figure 2 and the complete list of species is provided in Supplementary Table 2.

In total, 234 (91.0%) of the identified fungal species had never been associated with traditional agave fermentations before (Colón-González M., *et al*., 2024). From the core species, *P. polonicum*, *M. tassiana* and *A. pullulans* had not been reported previously. On the other hand, 23 species previously isolated using culture-based methods from traditional agave fermentations were not identified in our survey, although eleven of these are not included in the ITS database used here. At the level of genus, 90.7% of the 184 genera identified here had not been associated with traditional agave fermentations and we only missed three previously isolated genera (15%), all Basidiomycetes, and one of them is not included in the ITS database that we employed. Together, these results show that our effort greatly enriches our understanding of the fungal communities of these fermentations, adding a considerable number of species and genera that had not been associated with this environment.

### 3.4 Microbial diversity is only explained at the distillery level

To identify the geographic, climatic and production variables that influence the microbial composition of the fermentation tanks we analyzed the beta diversity generating dendrograms based on the similarities of the microbiomes of the samples (Figure 3). We employed an unweighted UniFrac distance matrix for bacteria and a Jaccard similarity matrix for fungi. The most important variables associated with each sample can be observed in Figure 3 as color bars beside the dendrograms. Surprisingly, the only grouping in which both fungal and bacterial ordination was statistically significant (ANOSIM) was according to the distillery (*r* = 0.5967, *P* = 0.0001 bacteria; *r* = 0.1703, *P* = 0.0315 fungi). This can also be appreciated at the dendrograms, where samples belonging to the same distillery often appear in the same clusters (Figure 3). No other variables were statistically significant for fungi, while grouping by fermentation stage, climate and region were also significant for bacteria (*r* = 0.2567, *P* = 0.006; *r* = 0.2768, *P* = 0.0001 and *r* = 0.1949, *P* = 0.0111, respectively).

**Figure 3.**
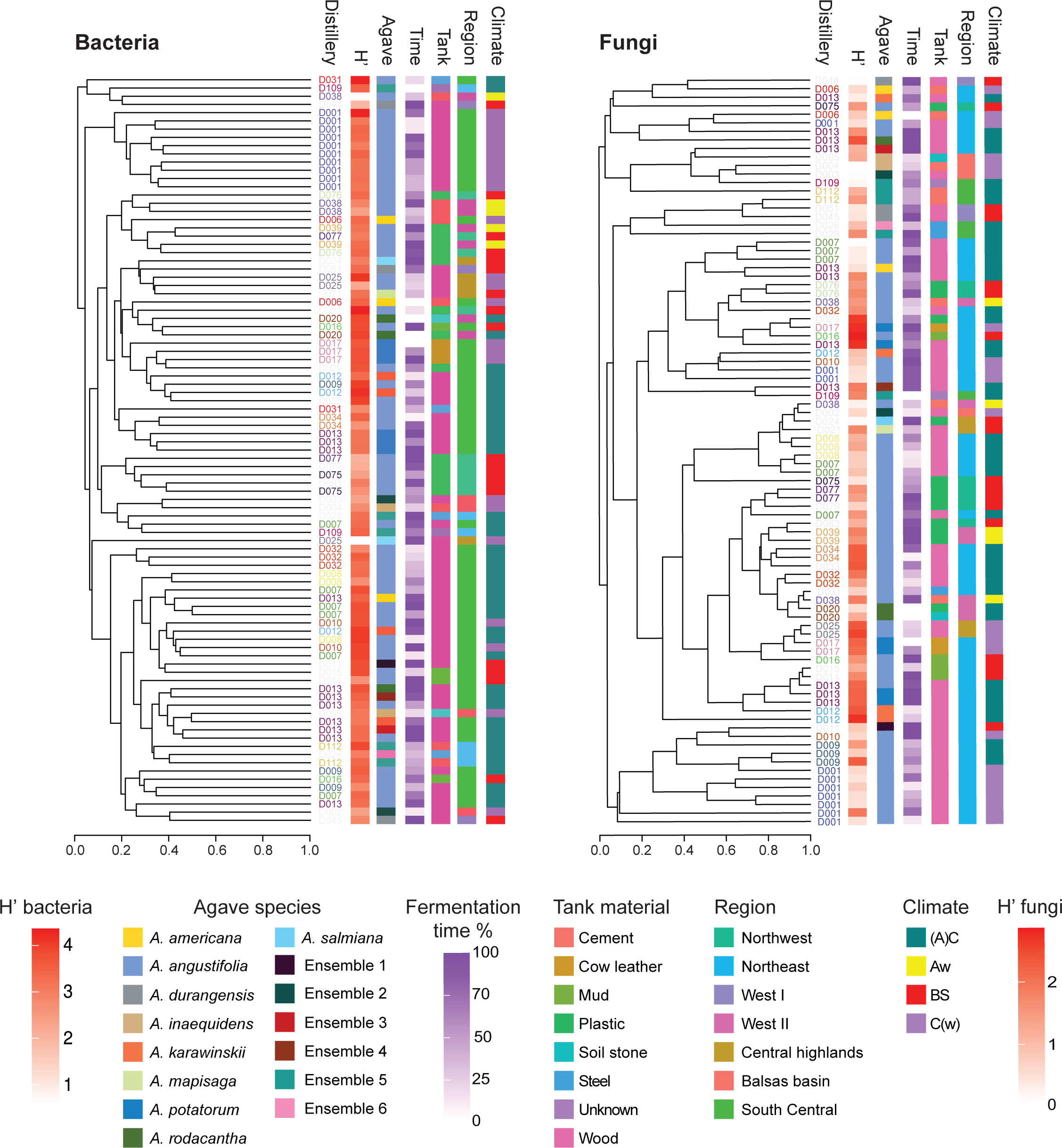
Analysis of the microbial diversity of agave fermentations and its relationship with the production practices and geographical characteristics. Dendrograms show the similarity of the bacterial (left panel) and fungal (right panel) microbiomes of the fermentations sampled. The same font color is used for samples from the same distillery. UniFrac distance was employed for bacteria while Jaccard similarity for fungi. The most important production and geo-climatic characteristics associated with each sample are shown in colored vertical bars at the right of each dendrogram. From left to right, Shannon diversity index (H’), agave species employed (Agave), fermentation stage expressed as percentage of the total fermentation time (Time), material of the tank (Tank), producing region as described in Figure 1 (Region) and climate group (Climate) are shown. The composition of fermentations with mixtures of agave plants (ensembles) is described in Supplementary Table 1. Climate groups according to Köppen and modified by E. Garcia (2004) were used. The four climate groups encompassing distilleries are tropical [Aw], subtropical [(A)C], semiarid [BS], and temperate [C(w)].

Given the clustering observed by distillery, we performed a Mantel test to evaluate if there is a correlation between geographic distance and microbial diversity of the fermentations. Using an unweighted UniFrac distance matrix for bacteria, we obtained an *r* value of 0.2321 with a significance of 0.001. The *r* and significance values using a Jaccard similarity matrix for fungi were 0.08247 and 0.101, respectively. For bacteria, these results showed that the microbial composition of two distilleries that are close by is more similar than the communities from distilleries that are further apart, which is in agreement with the ANOSIM results when assessing samples by distillery and region. In contrast, for fungi, the effect that we observed by distillery (ANOSIM) was not seen at all spatial scales. In summary, we observed that the distillery is an important determinant of both bacterial and fungal communities of the fermentation tanks, and, for bacteria, the fermentation phase, climate and producing region were also general determining factors.

### 3.5 Different bacterial and fungal community dynamics during *Agave* fermentation

Our general analyses of all distilleries revealed a significant association between the fermentation stage and the bacterial composition of agave fermentations. However, this was not observed for fungi, which was to some extent surprising given the ecological successions that have been observed in open fermentations employed for the production of other beverages (Pinto et al. 2015, Boynton and Greig 2016, Liu et al. 2020, Martiniuk et al. 2023). It is important to point out that the previous ANOSIM analysis was performed with the fermentation times of all sampled tanks. To investigate this further, we focused on two specific distilleries for which we had samples at different fermentation times (Figure 4). The samples of the two distilleries were grouped in three relative fermentation stages, each one representing a tercile of the total fermentation time (Initial, Mid and Final), and changes in diversity were evaluated through alpha diversity measures. The observed trends towards decreased diversity of both bacterial and fungal OTUs at the end of the fermentation were not statistically significant (Figure 4 and Supplementary Figure 1). However, for bacteria, the community structure was influenced greatly by the fermentation stage as reflected by the ANOSIM results when considering these two distilleries (*r* = 0.4608, *P* = 0.0128). In contrast, ANOSIM showed that the fungal communities are more uniform across the fermentation process (*r* = 0.02222, *P* =0.4398) as had been suggested by the analysis of all fermentation samples.

**Figure 4.**
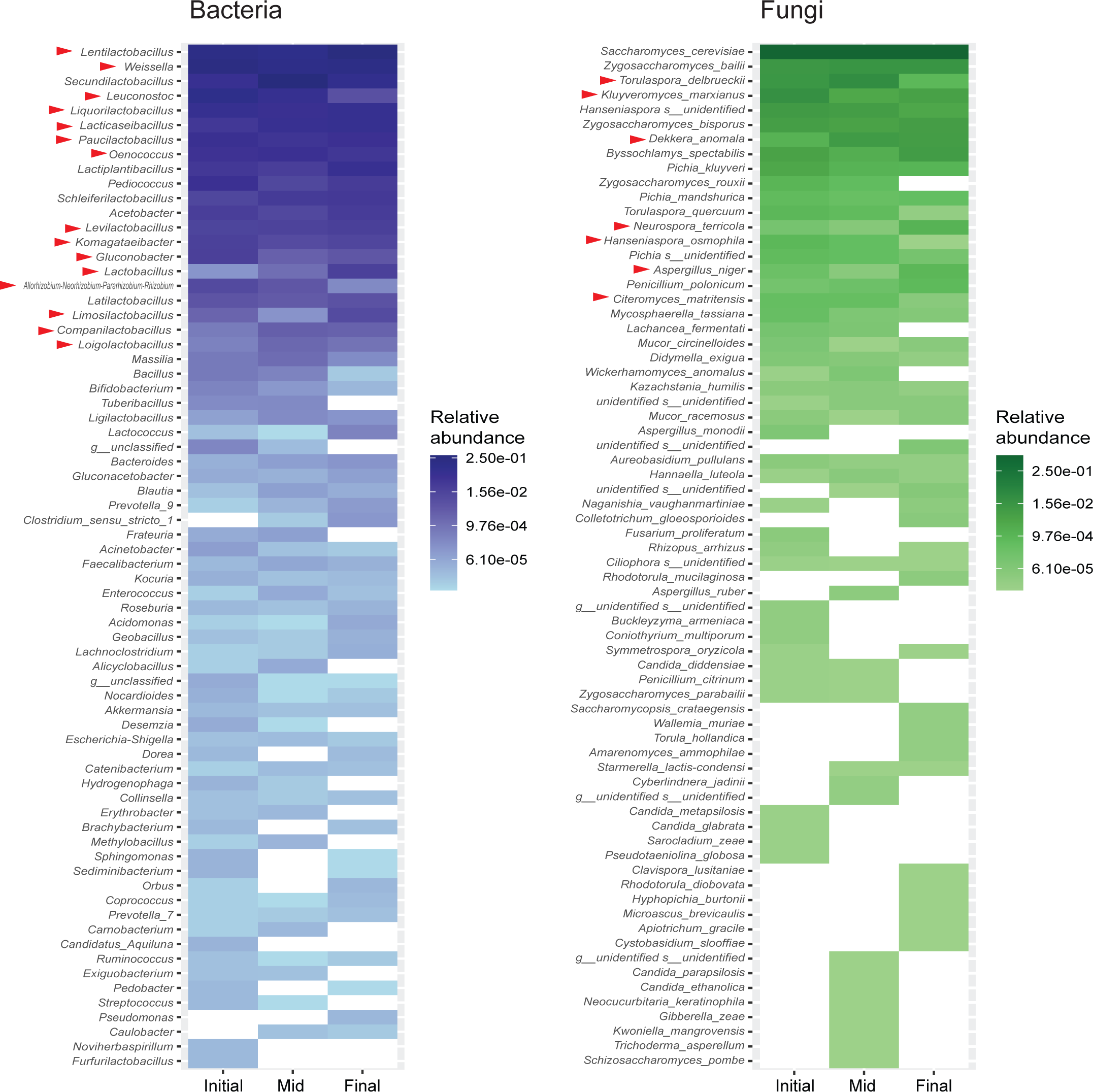
Microbial community changes through time during traditional agave fermentation. Relative abundance of the bacterial genera (left panel) and fungal species (right panel) identified in each of the three fermentation phases in two distilleries that had fermentations of different times when sampled. Only the 100 most abundant taxa are shown ordered from top to bottom according to their overall relative abundance. Red arrowheads denote taxons that were differentially enriched by DeSeq analysis as detailed in Supplementary Figure 2.

To determine if there were specific bacterial and fungal taxons enriched at the different fermentation stages, we performed DESeq analysis. For bacteria, we only found enriched genera when comparing the initial or mid stages with the final one, and there were more than double the number of genera enriched in the initial or mid stages (Supplementary Figure 2). *Allorhizobium, Komagataeibacter, Lentilactobacillus, Leuconostoc, Levilactobacillus, Liquorilactobacillus, Oenococcus,* and *Paucilactobacillu*s were enriched in both the initial and mid fermentation phases. The first two genera belong to the Proteobacteria and the rest are Firmicutes. *Companilactobacillus, Lactobacillus* and *Limosilactobacillus*, all Firmicutes, were enriched in the final stage compared to the two other stages. All the differentially enriched genera are part of the core bacterial component of agave fermentations.

In the case of fungi, there were only enriched species when comparing the initial and final fermentation stages with the mid phase (Supplementary Figure 2). *K. marxianus* was enriched at the beginning of the fermentation when compared to the mid stage, while *Dekkera anomala* showed the inverse enrichment. *Citeromyces matritensis, Hanseniaspora osmophila,* and *T. delbrueckii* were enriched in the mid fermentation phase when compared to the final stage, while *Aspergillus niger* and *Neurospora terricola* showed the reverse pattern. All these species are ascomycetes and *K. marxianus* and *T. delbrueckii* belong to the core fungal component of agave fermentations.

The statistically significant association between bacterial composition and fermentation stage suggested the possible occurrence of an ecological succession throughout agave fermentation, aligning with findings from other fermentation processes (Pinto et al. 2015, Boynton and Greig 2016, Liu et al. 2020, Martiniuk et al. 2023). However, in agave fermentations, the changes in fungal composition may be more subtle compared to other systems. Therefore, further analyses with increased sample size and specifically tailored to compare different stages are probably needed to detect these subtle variations in the fungal composition of these fermentations.

### 3.5 Specific bacterial genera are enriched in fermentations of certain agaves within a distillery

Although clustering by the agave species used in the fermentation was not statistically significant when considering all the samples, we included a distillery where fermentations of different agaves were being carried out in parallel (D13). This offered a unique opportunity to assess the influence of the plant employed as substrate because all other variables were similar between the tanks. As can be observed in the dendrograms of Figure 3, the bacterial microbiomes of the samples of *Agave potatorum* from this distillery clustered together while the ones of other agaves, including several of *A. angustifoli*a, did not. We did not observe clear clustering by agave species for the fungal microbiomes of this distillery. Performing a DeSeq analysis we identified 16 bacterial OTUs that were differentially enriched between the microbiomes of the *A. potatorum* samples and the rest of the agaves (Supplementary Figure 3). All of them but one are Firmicutes, and several belong to the core bacterial microbiome. Importantly, when the same analysis was performed, but instead comparing the *A. angustifoli*a fermentations against the rest of the samples, only one OTU was differentially enriched. Although further experiments are needed to better understand the influence of the agave species on the microbiome, our findings suggest that the specific biochemical composition of the plants could have determining effects on the microbial composition of agave fermentations, at least in the case of *A. potatorum*.

### 3.6 The microbiome of agave spirit fermentations and pulque are considerably different

Pulque is a traditional Mexican beverage produced from the fermentation of agave sap. Contrary to what is done to produce agave spirits, agave sap is not cooked for pulque production, and the beverage is the direct ferment without distillation. The microbiome involved in pulque fermentations has been well characterized using metagenomic approaches (Rocha-Arriaga et al. 2020, Astudillo-Melgar et al. 2023) and given the similarity in fermentation substrate, agave sap vs. cooked agave must, it is an excellent point of comparison to better understand the microbial communities involved in the production of agave spirits. For the analysis of pulque, we used 1,399,910 previously sequenced 16S and ITS pair-end reads (Rocha-Arriaga et al. 2020). These sequences were generated from six pulque samples with a mean of 74,882.16 ± 336.61 reads per sample for bacteria and 10,782.16 ± 280.71 reads for fungi. On average, the bacterial communities of pulque samples had a higher number of observed OTUs (3,105.5 ± 277.09) than samples from agave spirits (1,002.08). In agreement, the bacterial communities from agave spirit fermentations are less diverse than pulque, displaying a Shannon index (H’) of 3.26 ± 0.66, while the community of pulque showed 3.64 ± 0.51. Species dominance, evaluated through the Simpson index, presented similar values in agave spirit fermentations (0.86 ± 0.12) than in pulque samples (D = 0.85 ± 0.06). Similarly, the pulque microbiomes in average showed a higher amount of observed ITS OTUs (114.33 ± 98.82) than the ones from agave spirit samples (92.85 ± 39.5). For fungi both Shannon and Simpson indices were also more elevated for the communities of pulque samples (H’ = 2.18 ± 0.21; D = 0.80 ± 0.03 in pulque, against H’ = 1.20 ± 0.74; 0.44 ± 0.27 in agave spirits).

Overall, a total of 100 bacterial and 49 fungal genera were shared between pulque and agave spirit samples. Despite the communalities, the microbiomes of the two types of fermentations are clearly different. Agave spirit fermentations have 1,115 bacterial genera and 322 fungal species that are exclusive to these fermentations, while pulque fermentations have only 37 and 46, respectively. Furthermore, both bacterial and fungal beta diversity analyses by CAP showed evident separation of pulque and agave spirit microbiomes (Figure 5). Overall, each bacterial and fungal ordination explains 19 and 40% of the observed variance for bacteria and fungi, respectively, and showed that the communities from agave spirit fermentations differ more than the pulque samples (Figure 5). DESeq2 analysis identified a total of 31 bacterial genera that are overrepresented in agave spirit fermentations, and only eight that are overrepresented in pulque (Supplementary Table 3). Similarly, twelve species of fungi are significantly enriched in agave spirit fermentations, while eight species are overrepresented in pulque (Supplementary Table 3). In sum, our comparison showed important differences between the microbiomes of agave spirit fermentations and pulque even though both employ agave as the raw material.

**Figure 5.**
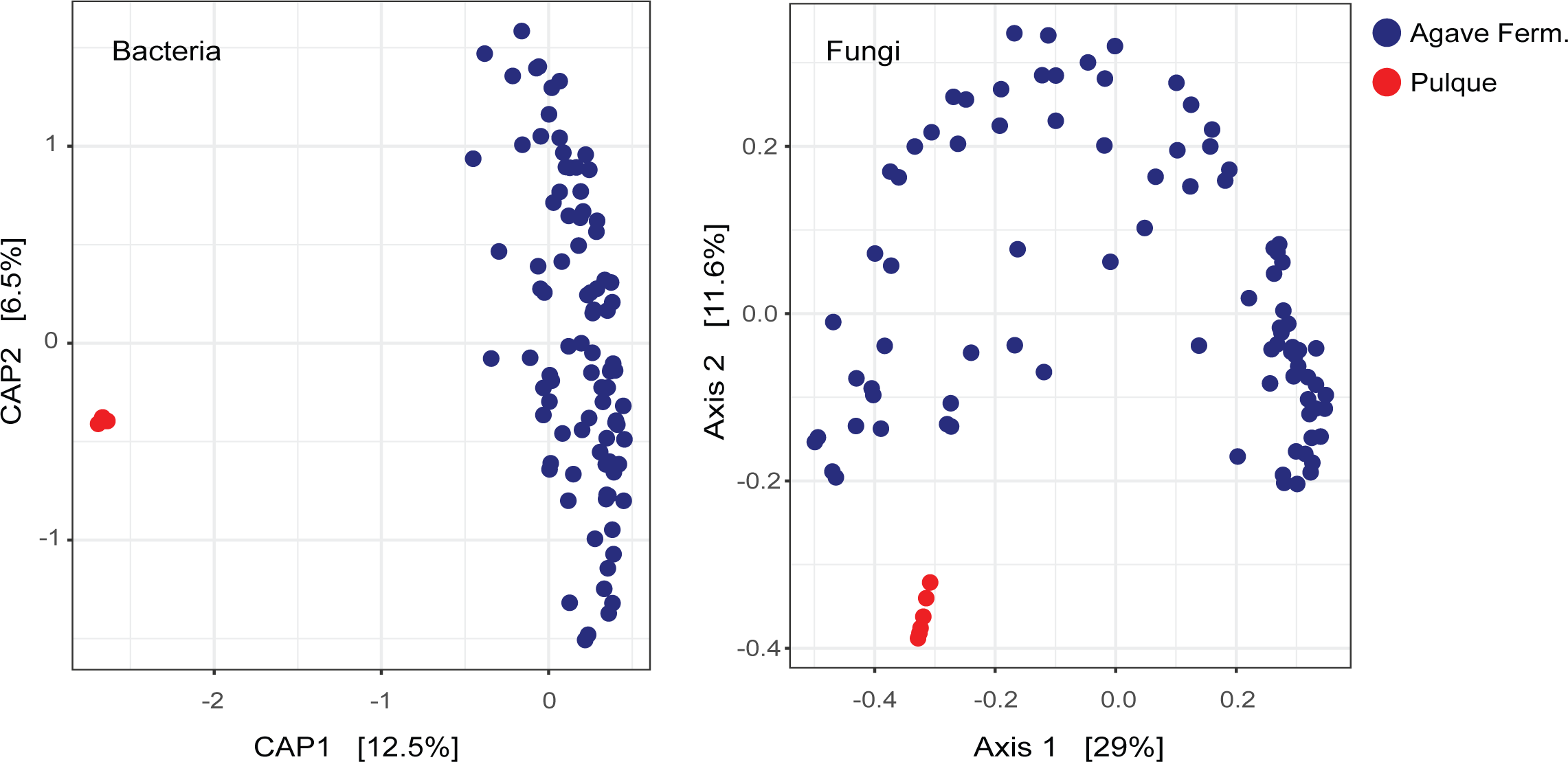
The microbiomes of pulque and agave spirit fermentations are considerably different. Beta diversity analysis of bacterial (left panel) and fungal communities (right panel) in pulque (red dots) and fermentations used for agave spirit production (blue dots). For bacteria we used PCA with an unweighted UniFrac distance matrix, and for fungi PcoA based on a Jaccard similarity matrix.

## 4 DISCUSSION

### 4.1 A country-wide analysis of microbial diversity in traditional agave fermentations

The production of agave spirits is culturally and commercially of central importance to Mexico. In recent decades, the aromas and flavors of these beverages are being increasingly appreciated worldwide, to the extent that other countries have also started producing agave spirits. Most of the focus on this activity has been placed on the plant variety and distillation practices employed, leaving aside the microorganisms that perform the fermentation (Arellano-Plaza et al. 2022). For example, the different designations of origin barely include this fundamental step. The microorganisms that perform the fermentation are particularly unknown for traditional distilleries in which no inocula are employed and the process is presumably performed by autochthonous bacteria and yeasts from the surrounding environment. To better understand these microbial communities, here we characterized the fungal and prokaryotic composition of a countrywide collection of fermentation musts employing a metabarcoding approach. Despite the wide range of geoclimatic characteristics and production practices encompassed by the distilleries, we did observe a core set of bacterial genera and fungal species that are prevalent in most fermentations (Table 1). Lactic acid bacteria were the most common, although we also identified a considerable number of acetic acid bacteria. In the case of fungi, Ascomycetes were by far the most ubiquitous and abundant since other phyla represented less than 10% of the total number of OTUs. These findings are in agreement with the microorganism that had been isolated in previous studies from this type of fermentations, which as a whole covered close to fifteen distilleries (Colón-González M., *et al*., 2024). Importantly, our study revealed extensive intra-generic diversity among prokaryotes and intra-specific variability among fungi, with the most abundant genera and species encompassing multiple OTUs.

Our work considerably expanded the list of microorganisms associated with traditional agave fermentations. This is especially true for prokaryotes that had been barely considered before – close to 98% of the bacterial genera that we identified had not been described in this environment. For fungi, over 90% of the genera and species detected had also not been isolated from these fermentations. Surprisingly, several of the taxa that we considered here as the core of these microbiomes (Table 1), had not been isolated in previous work, showing the importance of our effort to better understand these communities. For prokaryotes, this could be expected since only a few previous studies focused on these microorganisms (Escalante-Minakata et al. 2008, Narvaez-Zapata et al. 2010, Kirchmayr et al. 2017). In the case of fungi, it was surprising to find *P. polonicum*, *M. tassiana* and *A. pullulans* as prevalent taxa in agave fermentations. All these species are known to be widespread in a variety of environments and *P. polonicum* and *A. pullulans* have been used for biotechnological applications (Chi et al. 2009, Ding et al. 2013). On the other hand, *P. polonicum* is usually associated with food spoilage and *M. tassiana* belongs to a large genus of phytopathogenic fungi (Barr 1958, Duduk et al. 2014). It will be interesting to define whether these species contribute positively or negatively to the fermentation dynamics and organoleptic properties in agave spirit production.

Despite the breadth of our survey, we did miss bacterial and fungal genera that had been identified by previous studies. This agrees with our diversity estimations which suggested that a fraction of the microbial communities in each distillery was still to be determined. The unidentified microorganisms are most probably rarer species with focalized distribution since they were not identified in more than one previous study. Changes in the taxonomy of microorganisms also explained some of the discrepancies with previous reports as was the case for *Zymomonas*. Comparing the microbial diversity of agave fermentations with that of other fermented foods and beverages, the bacterial diversity (Shannon index) is close to average diversity in other fermentations, while fungal diversity is closer to the least diverse environments such as wine fermentations (Leech et al. 2020, Rocha-Arriaga et al. 2020, Kharnaior and Tamang 2023, Martiniuk et al. 2023, Qi et al. 2023). Overall, our work sets the basis for a more comprehensive understanding of the composition and diversity of the microbial communities responsible for the fermentation needed to produce traditional agave spirits.

### 4.2 Composition of microbial communities in agave fermentations is defined at the level of distillery

As in other open fermentation systems, it is expected that the climatic factors and elaboration practices will determine the microbial composition of agave fermentations at different geographical scales. For example, fermentations within a producing region would be expected to have microbiomes that are more similar to each other than to those of a different region. Our work offered the first opportunity to investigate such effects throughout the country as previous efforts had mainly focused on one or two distilleries and employed isolation methods that made it difficult to directly compare across studies. Surprisingly, we found that differences in the microbiomes were only observed for fungi and bacteria when comparing at the distillery level. This suggests that the microbiome of agave fermentations is firstly defined locally by the specific characteristics and practices of each production site. The few differences that we found in the microbiomes of tanks of distinct fermentation stages and agave plants within single distilleries is in agreement with this suggestion; the conditions within the distillery are more determinant than other factors.

Our results also showed that the fungal and bacterial components of agave fermentations are differentially influenced by environmental factors. Apart from the observations at the distillery level, significant differences were observed in the bacterial microbiomes of samples of distinct fermentation stages, climates and producing regions, but not in the fungal component of agave fermentations. Overall, we observed less diversity in the fungal microbiome with 13 species that form a core fungal component that is present in almost all fermentations sampled throughout the country (Table 1), and these species showed considerable intraspecific diversity. It remains to be seen whether fungal diversity within species is better correlated with geographical distribution.

Our findings are to some extent contrasting to what has been observed for microbiomes of wine fermentations, specifically for the fungal microbiome. The bacterial and fungal communities of wineries in California and Portugal, for example, have been reported to be specific to wine appellations, especially at early stages of fermentation (Pinto et al. 2015, Bokulich et al. 2016). It is worth pointing out that these regions are considerably smaller than the area where the agave fermentations sampled in this study are located and that endogenous microorganisms from the plant are not eliminated through a cooking step. Differences between the microbiomes of individual vineyards have also been found in these regions, similarly to what we observed between agave distilleries, but in addition there were differences between viticultural regions. These differences have been suggested to contribute to the specific characteristics of the wines from these appellations.

The wide range of geographical and climatic characteristics encompassed by the production sites of agave spirits, added to the great variety of cultural differences in production practices make agave fermentations a very diverse environment. Therefore, it is possible that a larger number of distilleries and fermentation tanks needs to be analyzed to better identify the factors that define the microbial communities of these fermentations. Yet, our work represents the most comprehensive effort to date to understand these microbiomes and provides valuable insights regarding the specific taxa that constitute them.

### 4.3 Fermentation time, agave species and production practices influence the microbiomes of agave fermentations

We observed a trend towards decreased alpha diversity of both bacterial and fungal OTUs towards the end of fermentation, even if the differences were not statistically significant (Supplementary Figure 1). The ANOSIM test revealed a significant influence of the fermentation stage on the structure of the prokaryotic communities in agave fermentations, highlighting a clear differentiation in bacterial beta diversity across stages. In addition, specific bacterial groups were enriched at different fermentation stages. The observed decrease in diversity towards the end of fermentation and the significant correlation between bacterial diversity and fermentation stage are consistent with findings from other fermentation processes (Pinto et al. 2015, Bokulich et al. 2016). In contrast, fungal communities appeared to be more uniform throughout the fermentations. In general, our results are in accordance with the survey of yeast diversity in traditional agave fermentations that we recently carried out employing culture-based methods (Gallegos-Casillas et al. 2023). This work centered in yeast communities, revealed composition changes through time, even if the alpha diversity was not statistically different across stages. However, in contrast with the metabarcodig results presented here, the previous analyses of yeast beta diversity did show a statistically significant decrease as a function of fermentation phase, even when the magnitude of the change was small (Gallegos-Casillas et al. 2023).

The observed decrease in diversity towards the end of fermentation and the statistically significant correlation between bacterial diversity and fermentation stage align with findings from other alcoholic fermentation processes (Costa et al. 2015, Pinto et al. 2015, Huang et al. 2023). However, in agave fermentations, the changes in fungal composition may be more subtle compared to other fermentations (Gallegos-Casillas et al. 2023). To gain a more comprehensive understanding of the ecological succession in spontaneous agave fermentations, further analyses with increased sample size and specifically tailored to compare different stages are necessary. These additional investigations will uncover finer details of microbial dynamics and shed light on the intricate ecological processes occurring during agave fermentation.

The plant species used to produce agave spirits is thought to be one of the most influential factors contributing to the flavor and aroma of these beverages. There are spirits made from agave varieties, such as *A. potatorum*, that are highly valued by consumers and their prices are considerably higher. It is possible that some of the attributes contributed by the specific agave variety may come indirectly from the influence that the plant components have on the microbial community responsible for the fermentation. By analyzing fermentations of different agave plants within the same distillery, we did observe specific bacterial genera associated with fermentations of *A. potatorum.* Clearly, further work is needed including sampling strategies specifically designed to better understand this association, but our findings suggest the possibility that the effects of the plant variety on the organoleptic properties of the spirits may occur by determining the composition of the microbiomes of the fermentations.

Production practices are another determinant of the quality of agave spirits. For example, distillation in clay pots is used in some breweries because it is thought to add specific flavors, despite the fact that it is a less efficient distillation system. In terms of a practice that could affect the fermentation process, we did not observe any statistically significant differences in the microbial communities when samples were grouped by the material of the fermentation tank and the grinding method used. As for other variables not found to be associated with the microbial composition, it is possible that a larger set of samples is needed to detect such effects given the complexity and diversity of the system. We did observe an important difference between the bacteria and fungi of pulque and agave spirit fermentations. At a fundamental level, the raw material used to produce both beverages are the oligosaccharides of agave plants. However, in the production of agave spirits these compounds have been subject to high temperatures for considerable periods of time. Apart from breaking oligosaccharides and chemically transforming other components of the plant, this step presumably eliminates all microorganisms from the plant and its surroundings. Although other variables could also be contributing to the differences with pulque, our work suggested that cooking the agave steams has important implications for the composition of the microbial communities of these fermentations.

### 4.4 Concluding remarks

To the best of our knowledge, the work presented here represents the most comprehensive analysis of the bacterial and fungal communities involved in the open fermentations needed to produce traditional agave spirits. We identified hundreds of species that had not been associated with this environment before and detected local differences in the microbiomes of the sampled distilleries. These local differences suggest that the sum of historical and environmental factors as well as the production practices of each distillery determine the microbial composition of agave fermentations. In turn, the specific microbiome of each distillery may contribute to the properties of the corresponding agave spirit, adding to the *terroir* of these beverages. Given the rapid pace at which the production of agave spirits is changing due to the increasing demand, the microbial communities here identified will serve as the basis to better understand and preserve the fermentations in this unique traditional process.

## FIGURE LEGENDS

**Supplementary Figure 1. Diversity analysis of the different fermentation stages in two distilleries.** Diversity estimators of the three fermentation stages analyzed in two specific distilleries that had tanks of different times when sampling took place.

**Supplementary Figure 2. DeSeq analyses of the different fermentation stages in two distilleries.** DeSeq results of the comparison of the three different fermentation stages at the level of OTU of the same two distilleries. Only comparisons where there were differentially enriched OTUs are shown.

**Supplementary Figure 3. DeSeq analysis in different agave species within a single distillery**

**Supplementary Table 1. Characteristics of the fermentation tanks from where the samples were collected.**

**Supplementary Table 2. Bacterial and fungal taxa identified in traditional agave fermentations.** This table contains four Tabs, two for prokaryotes (16S) and two for fungi (ITS). For each type of microorganism there is a list of all the OTUs identified (suffix “_OTU”) and of all the genera/species identified (suffixes “_Genus” and “_Species”).

**Supplementary Table 3. Bacterial and fungal OTUs and taxons identified in pulque**

## DATA AVAILABILITY

All data have been deposited at NCBI under BioProject PRJNA1085712.

## ACKNOWLEDGEMENTS

We are indebted to all the producers of agave spirits in the states of Durango, Guanajuato, Michoacan, Oaxaca, Puebla, Tamaulipas, Jalisco and Sonora, who kindly participated in this work by providing access to samples and for sharing their knowledge. Details about the acknowledgements regarding help in field work and sampling of the YeastGenomesMx consortium can be found in (Gallegos-Casillas et al. 2023). We thank Susana Ruiz-Castro, Porfirio GallegosLJCasillas, J. Abraham AvelarLJRivas and Luis F. GarcíaLJOrtega for technical assistance.

## AUTHOR CONTRIBUTIONS

Conceptualization: LM, AD, AH-L, EM. Methodology: AJ-S, LDA, SIJ-S. Field work: LM, AD, AH-L, EM. Investigation, laboratory work: SIJ-S, AE-J, IB. Investigation, formal analysis: AJ-S, LDA, EM. Visualization: AJ-S, LDA, EM. Funding acquisition: LM, AD, AH-L, EM. Supervision: AH-L, EM. Writing, original draft: AJ-S, EM. Writing, review and editing: LDA, LM, AD, AH-L. All authors read and approved the final version of the manuscript.

## MANUSCRIPT VERSIONS

The first version of this manuscript was released as a pre-print at *bioRxiv* (Jara-Servin A. et al., 2024).

