## Supplementary figures and images for "Microbial communities thriving in agave fermentations are locally influenced across diverse biogeographic regions"

### Supplementary Figure 1

##### Supplementary Figure 1

###### Bacteria

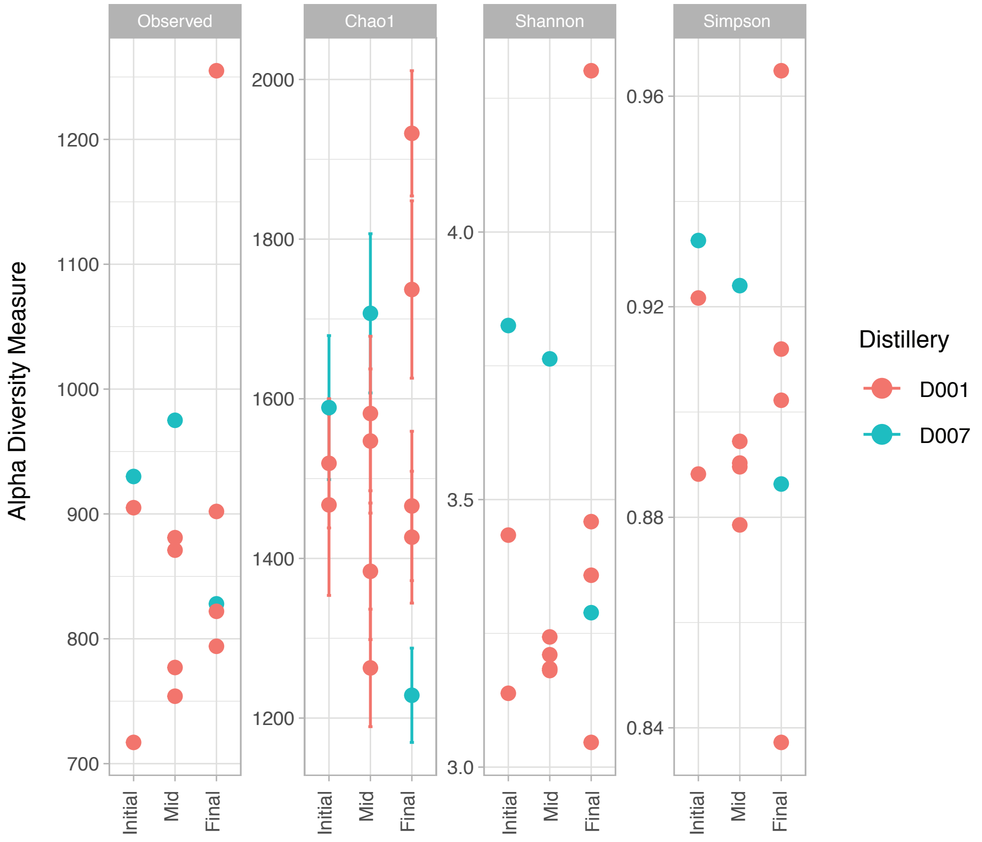


###### Fungi

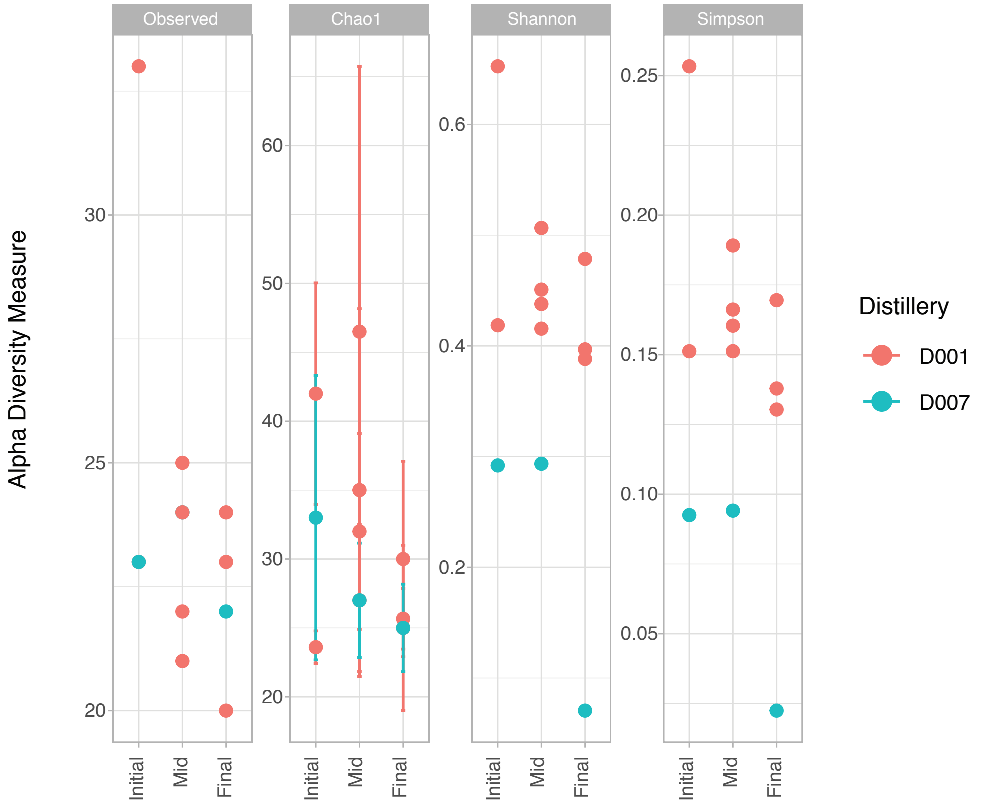

### Supplementary Figure 2

#### Supplementary Figure 2

##### Bacteria

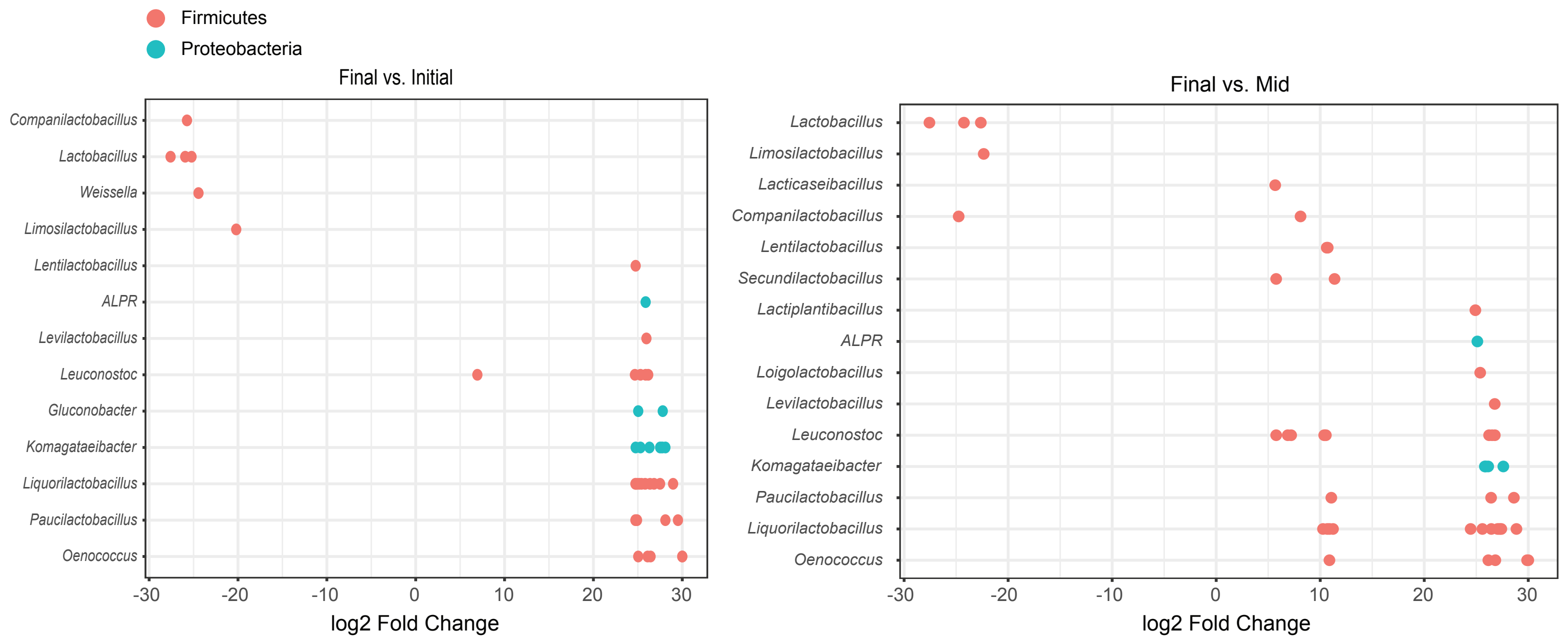


##### Fungi

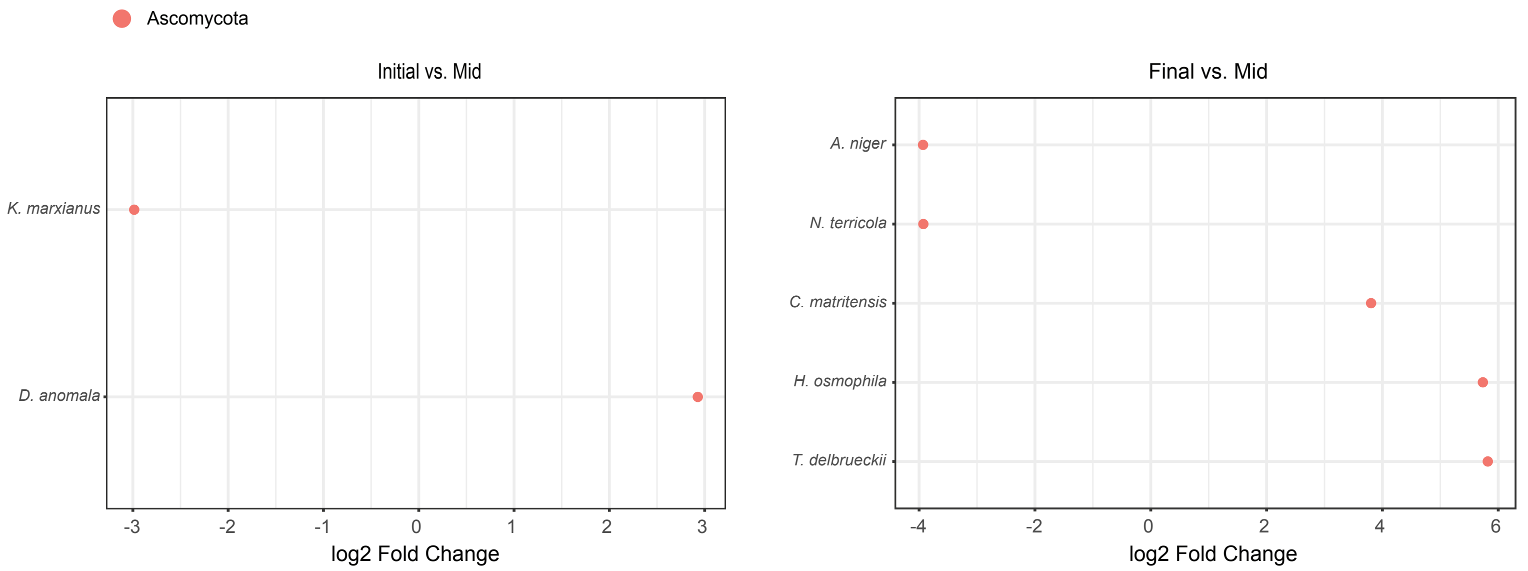

### Supplementary Figure 3

#### Supplementary Figure 3

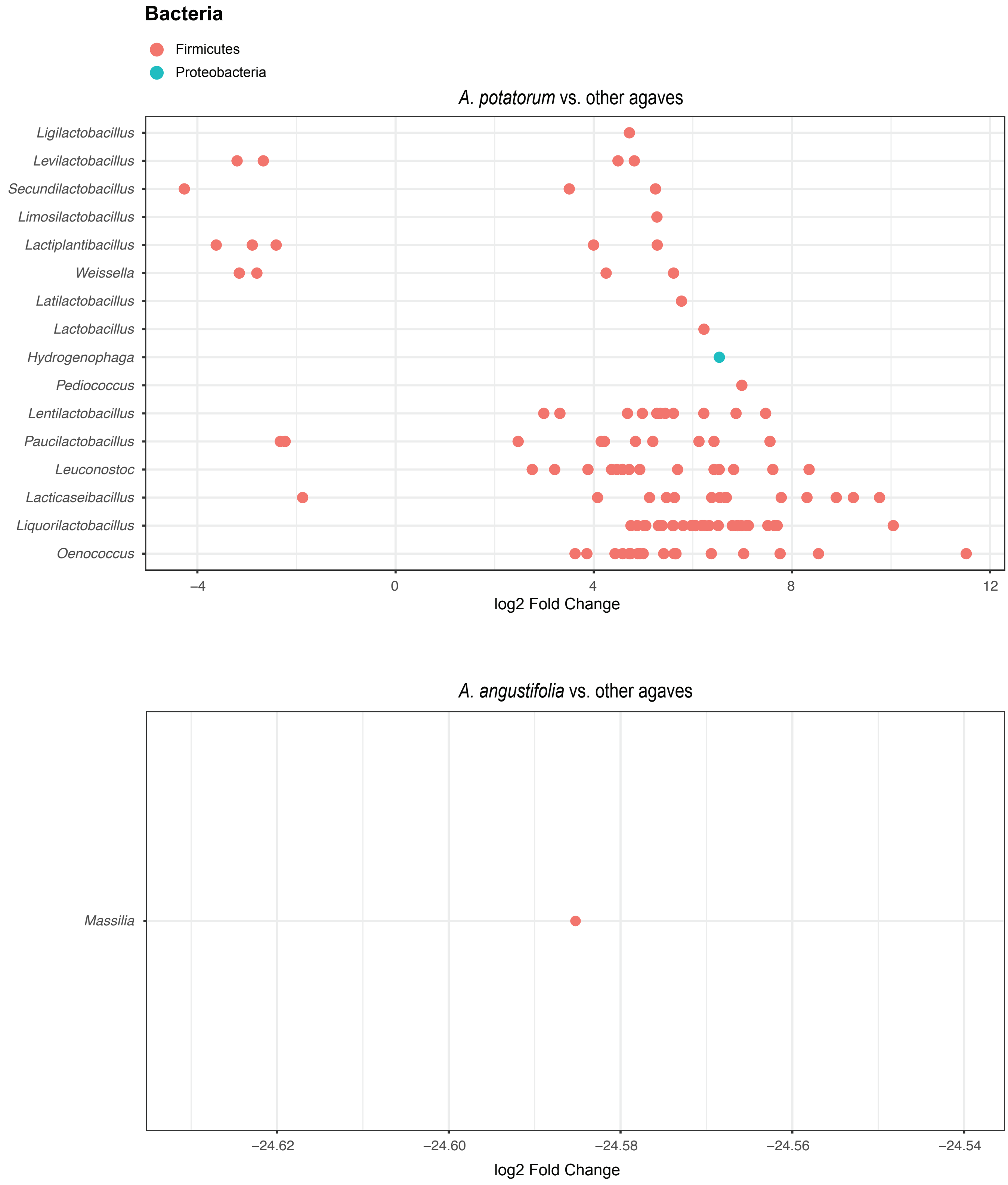
