## Supplementary Material 1 for "Microbial communities thriving in agave fermentations are locally influenced across diverse biogeographic regions"

**"Thriving Amidst Sweetness and Harshness: The Life of Yeasts in the Agave Fermentation Environment"**

Maritrini Colón-González[1,3], Xitlali Aguirre-Dugua[2], Mariana G. Guerrero-Osornio[1], J. Abraham Avelar-Rivas[3], Alexander DeLuna[3], Eugenio Mancera[4], Lucía Morales[1]

[1]*Laboratorio Internacional de Investigación sobre el Genoma Humano (LIIGH), Universidad Nacional Autónoma de México, 76230 Juriquilla, Mexico*

[2]*Investigadores por México, Consejo Nacional de Humanidades, Ciencias y Tecnologías (Conahcyt), 03940 Ciudad de Mexico, Mexico*

[3]*Unidad de Genómica Avanzada (Langebio), Centro de Investigación y de Estudios Avanzados del Instituto Politécnico Nacional, 36824 Irapuato, Mexico*

[4]*Departamento de Ingeniería Genética, Unidad Irapuato, Centro de Investigación y de Estudios Avanzados del Instituto Politécnico Nacional, 36824 Irapuato, Mexico*

**Keywords:** agave, fermentation, yeast, microbiome, domestication

**Running Title:**
Yeasts of the sweet and harsh agave environment

**Take Away:**
The overall purpose of this review is: i) to describe the environmental and productive context in which open agave fermentations occur, ii) to summarize the current knowledge about yeasts' traits that allow adaptation to this environment, and iii) to discuss how human practices may have inadvertently shaped the diversity of the agave fermentation microbial community.
